# Protein network structure enables switching between liquid and gel states

**DOI:** 10.1101/754952

**Authors:** Jeremy D. Schmit, Jill J. Bouchard, Erik W. Martin, Tanja Mittag

## Abstract

Biomolecular condensates are emerging as an important organizational principle within living cells. These condensed states are formed by phase separation, yet little is known about how material properties are encoded within the constituent molecules and how the specificity for being in different phases is established. Here we use analytic theory to explain the phase behavior of the cancer-related protein SPOP and its substrate DAXX. Binary mixtures of these molecules have a phase diagram that contains dilute liquid, dense liquid, and gel states. We show that these discrete phases appear due to a competition between SPOP-DAXX and DAXX-DAXX interactions. The stronger SPOP-DAXX interactions dominate at sub-stoichiometric DAXX concentrations leading to the formation of crosslinked gels. The theory shows that the driving force for gel formation is not the binding energy, but rather the entropy of distributing DAXX molecules on the binding sites. At high DAXX concentrations the SPOP-DAXX interactions saturate, which leads to the dissolution of the gel and the appearance of a liquid phase driven by weaker DAXX-DAXX interactions. This competition between interactions allows multiple dense phases to form in a narrow region of parameter space. We propose that the molecular architecture of phase-separating proteins governs the internal structure of dense phases, their material properties and their functions. Analytical theory can reveal these properties on the long length and time scales relevant to biomolecular condensates.

---

Many biomolecules have been shown to spontaneously form condensed states that serve organizational or functional roles in living cells [1, 2]. Often these condensates are responsive to changes in environmental conditions, cell cycle, or stress [3–6]. While many simple globular proteins undergo phase separation, their phase boundaries are usually in non-physiological, or even inaccessible, regions of phase space due to the weak intermolecular interactions [7, 8]. In contrast, multi-valent proteins have evolved to drive biologically relevant phase transitions; a higher valence means that a lower total protein concentration is sufficient to achieve phase separation [9]. Many proteins that drive phase separation are intrinsically disordered with attractive moieties separated by long flexible linkers. This has inspired polymer theory-based models to describe the effects of the attractive “stickers” [10] and the flexible, inert “spacers” [11] to describe the condensation of these molecules into a polymer fluid. However, there is considerable heterogeneity in the protein architectures that have evolved to form condensates. Phase separation can be driven by discrete folded domains that specifically recognize linear motifs in binding partners [9] and the linkers between domains/motifs can be long, short, or nonexistent. In other cases the driving force is weak contacts between sidechains or backbones of disordered regions [5, 10, 12]. The existence of these different modes of interactions is thought to provide the specificity to facilitate the coexistence of multiple different biomolecular condensates and the conversion between different condensed states.

In this paper we study phase transitions involving the speckle-type POZ protein (SPOP). SPOP is the substrate adaptor of a ubiquitin ligase [13, 14] and recruits substrates such as the androgen receptor (AR) [15], bromodomain-containing proteins (BRD2, BRD3 and BRD4) [16, 17], and the death-domain-associated protein (DAXX) [18] for ubiquitination and subsequent degradation. SPOP is the most frequently mutated gene in prostate cancer and the cancer-associated mutations affect the interaction between SPOP and these substrates [19–22]. We recently showed that SPOP can undergo phase separation with cDAXX, the intrinsically disordered C-terminal tail of DAXX, and that their phase behavior is complex [23]. In isolation, SPOP polymerizes into linear assemblies [24], while cDAXX forms liquid droplets. When small amounts of DAXX are added to SPOP solutions the result is a crosslinked gel, indicative of a kinetically arrested state. However, the addition of more cDAXX (but less than the amount required to form cDAXX droplets) dissolves the gel, resulting in liquid droplets that contain both SPOP and cDAXX [23].

Protein condensates are disordered and, therefore, it is difficult to probe the internal structure by experiments. Furthermore, the dense phases form on timescales of seconds to hours, which presents a severe challenge for simulation-based approaches. Here we obtain structural insight by developing a theory that captures the phase boundaries and condensate densities. We show that the gel phase is not driven by the binding energy of DAXX crosslinks, but rather by the entropy of multivalent DAXX binding. We also show that the liquid-gel transition is caused by a competition between SPOP-DAXX and DAXX-DAXX interactions, with the former giving way to the latter as the binding sites on SPOP become saturated. As a result, the addition of more DAXX simultaneously lubricates the kinetically trapped gel and introduces the liquid state as the free energy minimum.

## I. RESULTS

### A. SPOP-SPOP interactions cause the formation of linear rods

SPOP is a three-domain protein, which binds substrates through the MATH domain (green in Fig. 1) and self-associates via two dimerization domains (red and blue in Fig. 1) [24–26]. The strong binding interface (BTB-BTB) leads to the formation of dimers with nanomolar affinity. Since the concentrations leading to dimers are considerably lower than the micromolar concentrations explored by Bouchard et al. [23], we neglect the presence of monomers and consider the assembly of a solution of dimers at concentration *c*_*s*_*/*2, via the BACK domain which has a micromolar affinity in the absence of crowding agents. Each dimer has two identical dimerization interfaces that can self-associate with an association constant *k*_*s*_ = *c*_*n*+2_*/*(*c*_*n*_*c*_2_), where *c*_*i*_ is the concentration of SPOP assemblies containing *I* monomers. Since these assemblies are linear and rigid on the lengthscale set by the DAXX binding sites, we refer to SPOP assemblies as rods throughout the paper. The concentration of SPOP rods of all lengths is

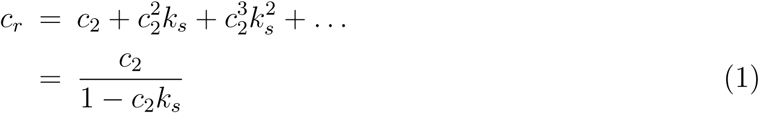

The total concentration of SPOP in the system is *c*_*s*_ = 2*c*_2_*∂c*_*r*_*/∂c*_2_, which can be inverted to give the free dimer concentration as a function of the total concentration

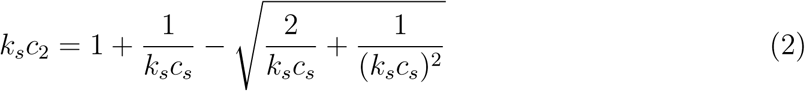

The average assembly length, measured in dimer units, is

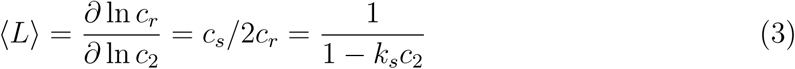

This function is plotted in Fig. 1 showing that the average length of SPOP assemblies ranges from 20-100 dimer units over the experimental concentration range [24].

**FIG. 1:**
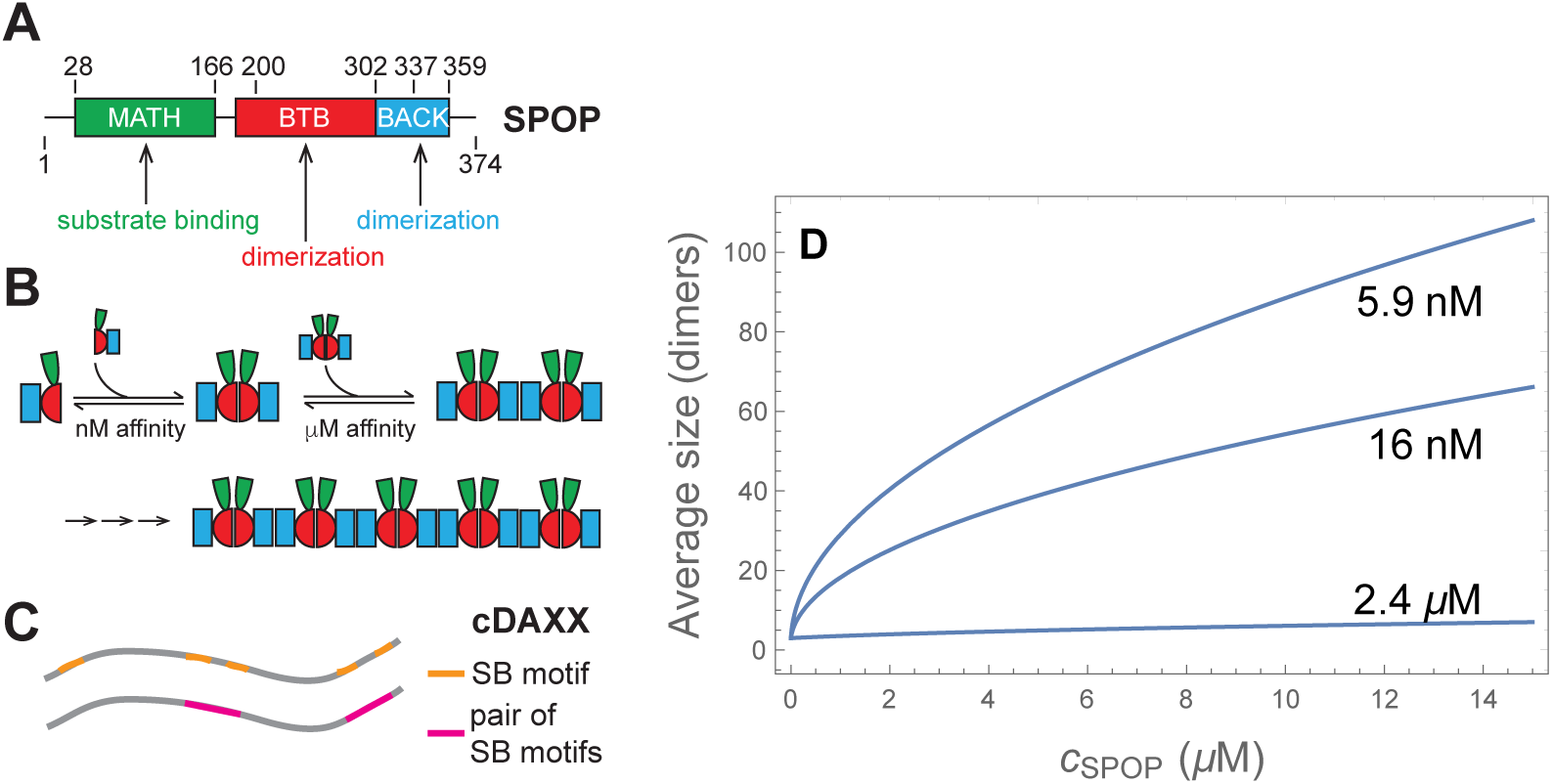
(A) The three domains of SPOP determine its assembly behavior. (B) BTB-BTB interactions lead to the formation of dimers with a nM affinity. BACK-BACK interactions allow for the polymerization of dimers to form linear rods. The MATH domain binds to substrates such as DAXX. (C) cDAXX, which is the intrinsically disordered C-terminus of DAXX and encompasses residues 495-740 of DAXX, has five SPOP binding (SB) motifs. Four of these are arranged in pairs that bind cooperatively to pairs of MATH domains. The remaining SB motif is weak and is neglected in our model. (D) Average length of SPOP assemblies (measured in dimer units) for three values of the polymerization constant, as calculated from Eq. 3. In the absence of ficoll the polymerization constant is 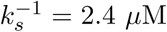. The other two curves represent the range of values consistent with the phase diagram observed in 4% ficoll.

### B. Entropy of SPOP-DAXX binding depends on the aggregation state

Since our theory is expected to apply equally to DAXX and cDAXX (DAXX^495−740^), in the remainder of the paper we will use the term DAXX unless specifically referring to cDAXX. DAXX has five appreciable SPOP recognition motifs that can bind to the MATH domain on a SPOP monomer [23]; four of these are located in pairs separated by 9 and 17 amino acids, respectively (Fig. 1C). Given the small separation, we assume that these pairs will bind cooperatively to both sites on a SPOP dimer giving DAXX an effective valence of two (neglecting the weak fifth binding motif). In the remainder of the paper, references to DAXX binding sites will refer to a pair of SPOP recognition motifs and a binding event will occupy both sites on a SPOP dimer.

We compute the free energy of a solution of *M* SPOP dimers. We divide these into three groups. *N*_0_ dimers are unbound, *N*_1_ are bound to a DAXX molecule that has a free second site, and 2*N*_2_ are bound to a DAXX molecule that binds a second SPOP dimer with its other binding site. Therefore *M* = *N*_0_ + *N*_1_ + 2*N*_2_ and the total number of DAXX molecules that are bound to SPOP is *N*_1_ + *N*_2_. The free energy of the system is

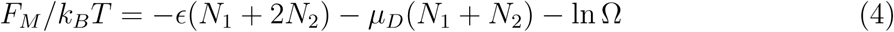

where *ϵ >* 0 is the free energy of a SPOP dimer binding to a DAXX binding site and *µ*_*D*_ is the chemical potential of DAXX (both scaled by *k*_*B*_*T* to make them dimensionless). Eq. 4 neglects DAXX-DAXX interactions. These can be included using a term proportional to (*N*_1_ +*N*_2_)^2^, which can be treated using a self-consistent mean-field approximation. However, this adds considerable complexity with minimal additional insight.

The difficulty in *F*_*M*_ is calculating Ω, which is the number of ways to arrange the DAXX molecules on the binding sites. To do this, we use a method inspired by the Flory-Huggins lattice model [27, 28], which yields (see Methods)

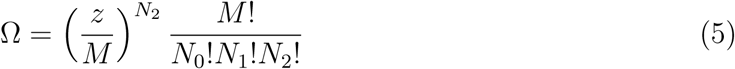

where *z* is a parameter describing the number of SPOP dimers within reach of a DAXX molecule bound with one site. Minimizing the free energy with respect to the site occupancies (see Methods) yields

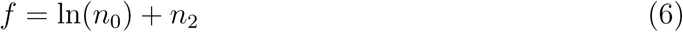

and the site occupancies

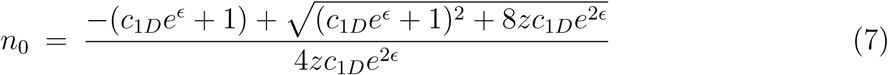

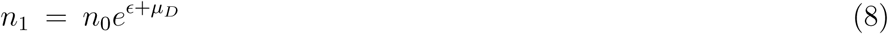

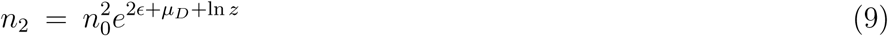

where *f* = *F*_*M*_ */k*_*B*_*T*, *n*_*i*_ = *N*_*i*_*/M*, and *c*_1*D*_ is the concentration of unbound DAXX.

Double bound DAXX molecules are of particular interest because these are the molecules that crosslink the rods into a gel. *n*_2_ is a non-monotonic function of the DAXX concentration (Fig. 2a). At low DAXX concentrations nearly all bound DAXX molecules bind with both sites. However, as the number of free binding sites *n*_0_ vanishes, only single bound molecules fit. This puts a limit on the concentration range where crosslinking can occur.

**FIG. 2:**
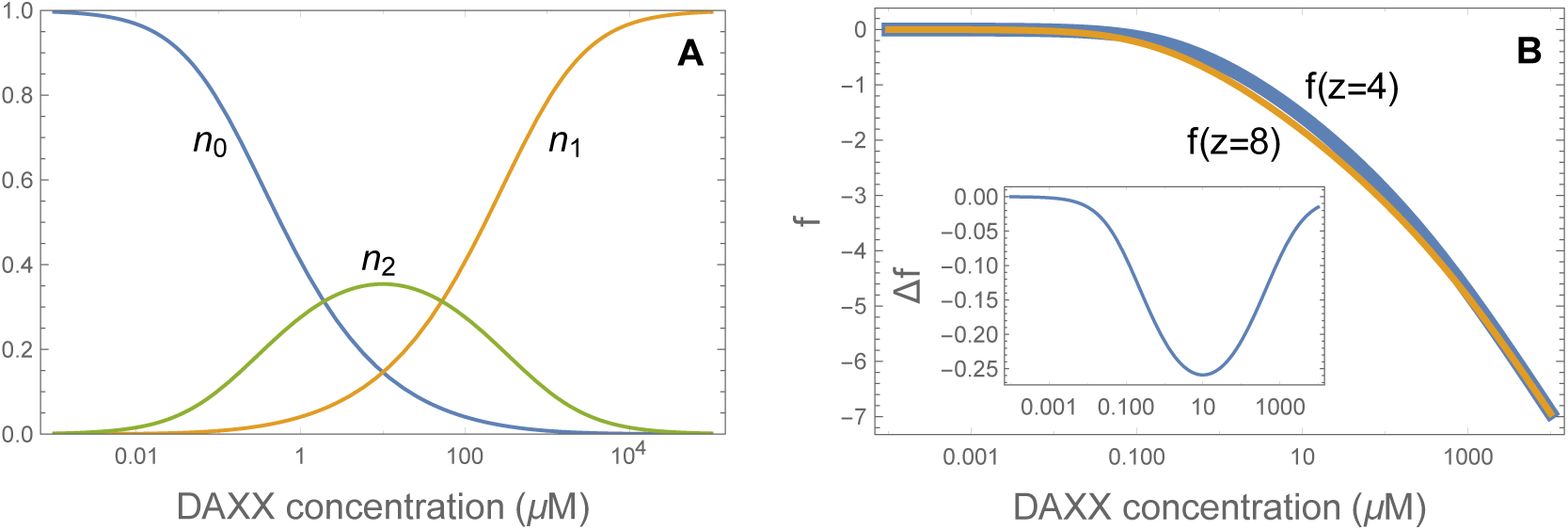
(A) Double bound DAXX molecules are only found in a limited concentration range, as seen by the non-monotonic behavior of *n*_2_. At low DAXX concentration there is an abundance of free binding sites and most DAXX molecules are double bound. However, as the number of binding sites saturates, it becomes progressively harder to find a nearby second binding site. Eventually, all binding sites are filled with single bound DAXX. (B) Comparison of the free energy for two different binding site densities. The high density system is lower in free energy for values of *c*_1*D*_ where DAXX is double bound. (inset) Free energy difference between the gel phase (*z* = 8) and the vapor phase (*z* = 4).

The free energy difference between the gel and soluble states arises from the parameter *z*, which determines the entropy of double bound DAXX. From Eq. 6, the free energy change from the increased entropy is

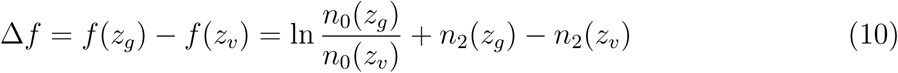

where *z*_*g*_ is the number of nearby binding sites in the gel state and *z*_*v*_ ≃ 2*R*_*g*_*/D*_*S*_ is the number of nearby sites for a SPOP rod in the vapor phase, where *R*_*g*_ is the radius of gyration of the disordered region between DAXX binding sites and *D*_*S*_ is the diameter of a SPOP dimer. The free energy difference (plotted in the inset of Fig. 2B) is small, on the order of 0.1 *k*_*B*_*T* per site. The contribution arising from the binding entropy is *n*_2_ ln(*z*_*g*_*/z*_*v*_), which accounts for nearly all of the free energy change. Δ*f* vanishes at low and high DAXX concentrations where *n*_2_ is small. This limited range of attraction allows for re-entrant behavior similar to that seen for RNA condensation due to charge-charge interactions [29].

### C. Favorable DAXX-DAXX interactions lead to a liquid state

At high concentration DAXX condenses into a liquid state. This condensation is driven by a favorable interaction per molecule that we denote *ϵ*_*D*_. When DAXX is mixed with SPOP, a rod of length *L* will acquire a layer of adsorbed DAXX. If we assume the double bound molecules are too constrained to participate in intermolecular interactions, *N*_*D*_ = *Ln*_1_ DAXX molecules are available for liquid contacts. When this assembly enters a DAXX fluid, the attractive energy will be approximately *N*_*D*_*ϵ*_*D*_. This additive approximation is justified by the similar concentrations of the DAXX fluid and the DAXX condensed on SPOP (see Section I F). This mechanism lowers the saturation concentration of the DAXX because SPOP scaffolds the formation of high molecular weight DAXX assemblies. Once again, this mechanism is not unique to the SPOP system as it has similarities to the condensation of RNA coated by intrinsically disordered proteins [30].

### D. Phase diagram can be computed using dilute solution affinity measurements

To compute the phase behavior of the system, we write down the chemical potentials for SPOP rods of length *L* (given by Eq. 3) in the vapor, liquid, and gel states

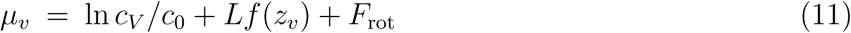

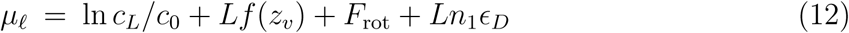

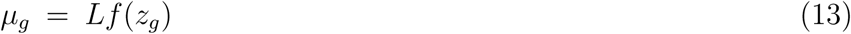

where *c*_*V*_ and *c*_*L*_ are the SPOP rod concentrations in the vapor and liquid phases, respectively. Each expression contains a term *Lf* accounting for the free energy of SPOP-DAXX binding. The vapor and liquid phases have terms for rotational entropy *F*_*rot*_, and translational entropy (scaled by a reference concentration *c*_0_). These contributions are neglected for the gel phase, which we expect to have solid-like ordering locally, even if it is disordered on longer length scales [23]. Finally, the expression for the liquid contains a term for DAXX-DAXX interactions. To mimic the crowded conditions of the cell, our experiments are conducted with added ficoll. This will enhance binding interactions (which is captured by the affinity parameters) as well as introduce non-specific depletion interactions. Eq. 13 neglects non-specific SPOP-SPOP interactions which can be added using a term proportional to *L* and the ficoll concentration. This omission results in minor qualitative shifts to the phase boundaries at 4% ficoll, but the depletion term is necessary to capture the larger changes at ∼ 10% ficoll.

**TABLE I:**
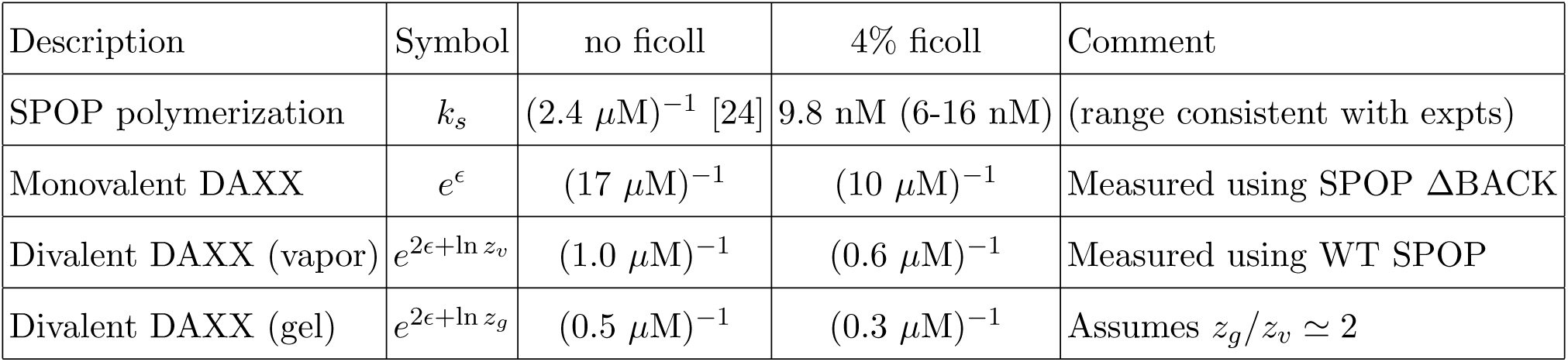
Association constants used in the calculations.

| Description | Symbol | no ficoll | 4% ficoll | Comment |
| --- | --- | --- | --- | --- |
| SPOP polymerization | $k_s$ | $(2.4 \mu\text{M})^{-1}$ [24] | 9.8 nM (6-16 nM) | (range consistent with expts) |
| Monovalent DAXX | $e^\epsilon$ | $(17 \mu\text{M})^{-1}$ | $(10 \mu\text{M})^{-1}$ | Measured using SPOP ΔBACK |
| Divalent DAXX (vapor) | $e^{2\epsilon+\ln z_v}$ | $(1.0 \mu\text{M})^{-1}$ | $(0.6 \mu\text{M})^{-1}$ | Measured using WT SPOP |
| Divalent DAXX (gel) | $e^{2\epsilon+\ln z_g}$ | $(0.5 \mu\text{M})^{-1}$ | $(0.3 \mu\text{M})^{-1}$ | Assumes $z_g/z_v \simeq 2$ |

We identify the quantity *k*_2_ = *e*^2*ϵ*+ln(*z*)^ as the association constant for divalent SPOP-DAXX binding. This is the reciprocal of the dissociation constants reported in Fig. 3 (in the vapor phase) so *k*_2*v*_ = (1.0 *µ*M)^−1^. In the gel phase we have 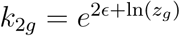 or 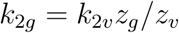. Similarly, we identify the quantity 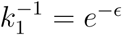 with the association constant for DAXX bound to a single SPOP dimer. This was measured by deleting the BACK domain so that SPOP assembly is arrested at the dimer. From these measurements we find *k*_1_ = (10 *µ*M)^−1^ in the presence of ficoll (Fig. 3). The SPOP polymerization affinity is 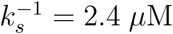 in the absence of ficoll [24]. The SPOP-SPOP interfaces exclude a larger volume to ficoll than the SPOP-DAXX interface, so it will be more strongly affected by ficoll. The latter affinity is enhanced by ln(17*µ*M*/*10*µ*M) *≃* 0.5*k*_*B*_*T* in the presence of ficoll. We find the best agreement with the experimental phase diagram when the SPOP-SPOP affinity is enhanced by 5-6 *k*_*B*_*T*, corresponding to 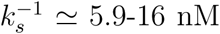. The other free parameters in the model are *z*_*g*_*/z*_*v*_ and *B*, which have the effect of expanding and contracting the gel phase, respectively. Fig. 4 shows the phase diagram for *z*_*g*_ = 2*z*_*v*_, *B* = 0.8 *µ*M, and 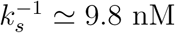 (equivalent to a 5.5 *k*_*B*_*T* ficoll effect).

**FIG. 3:**
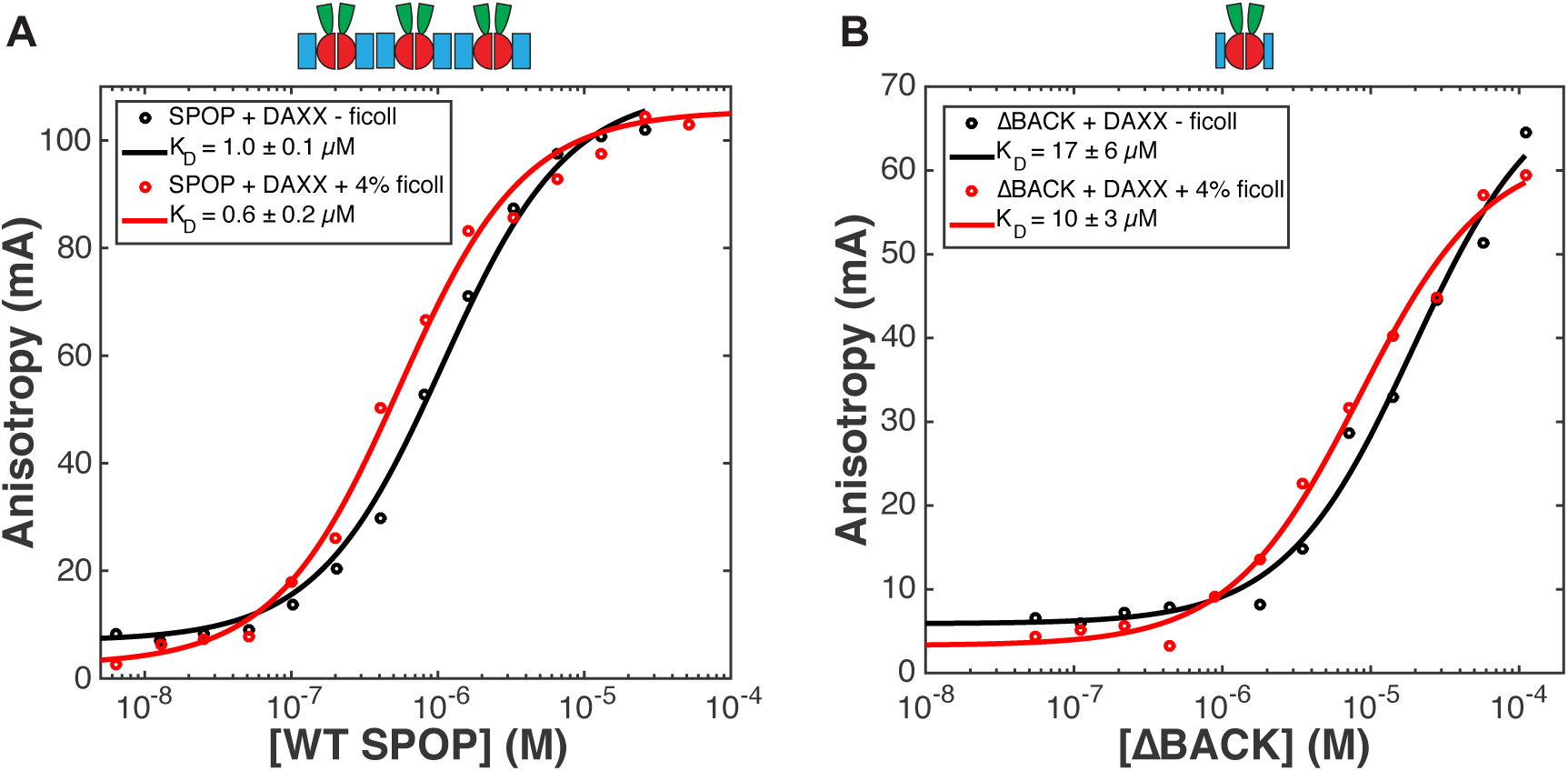
Binding curves in the presence and absence of ficoll. Representative fluorescence anisotropy direct binding curves of A) WT SPOP^28−359^ and B) SPOP ΔBACK^28−337^ from triplicate measurements with rhodamine-labeled His-cDAXX *±*4% ficoll. Experimental data points are shown as circles, and non-linear least squares fits are shown as lines. Average *K*_*D*_ values from triplicate experiments are reported in the legends, the errors represent standard deviations.

**FIG. 4:**
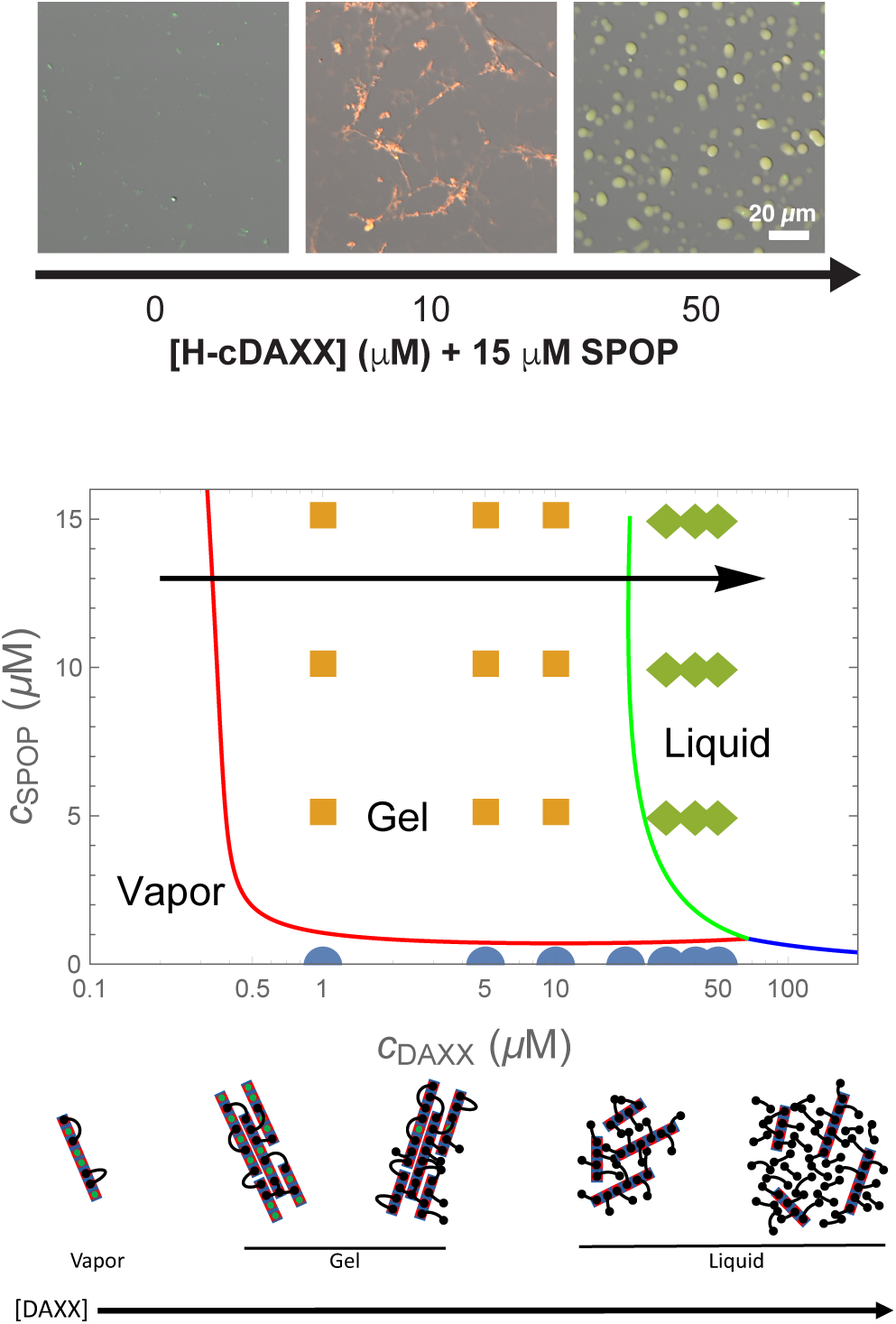
(top) Micrographs (fluorescence overlaid with DIC) of 15 *µ*M SPOP solutions with increasing concentrations of cDAXX showing the vapor, gel, and liquid phases in 4% ficoll. (middle) Phase diagram calculated from Eqs. 25, 26, and 28. Circles, squares, and diamonds indicate vapor, gel, and liquid phases observed experimentally [23]. The calculation shows the behavior of SPOP assemblies and does not account for the pure DAXX fluid. (bottom) Cartoon of the progression of structures along the arrow in the phase diagram. At low DAXX concentration there are insufficient crosslinks to drive condensation and the SPOP assemblies are in the vapor phase. The gel is formed when the excess entropy of arranging the DAXX molecules is enough to offset the translational entropy of condensation. When the binding sites become saturated, double bound DAXX molecules become rare, however, the SPOP assemblies are still held together in the liquid phase by weak DAXX-DAXX interactions. Finally, at very high concentrations, DAXX monomers condense into a liquid that dissolves the SPOP assemblies.

### E. Phase diagram shows a range of concentrations with a crosslinked gel state

The observed behavior is explained by the cartoons in the bottom panel of Fig. 4. At low DAXX concentrations the small enhancement in binding entropy is not sufficient to overcome the translational entropy loss of condensation, so the rods remain in the vapor phase. The gel appears when the increase in binding entropy is able to overcome this penalty. Although the benefit per molecule is small (Fig. 2B), the net gain can be substantial due to the large number of binding sites on the SPOP assemblies. Note that the high density of binding sites in the gel phase is possible because the rigid SPOP assemblies have a minimal conformational entropy penalty for stacking. This would not be the case for a molecule connected by flexible linkers instead of rigid polymerization interfaces.

As the SPOP sites saturate, the number of double bound DAXX molecules is insufficient to maintain the condensed state. This would lead to a return to the vapor state if not for the favorable DAXX-DAXX interaction. While these interactions are weaker than the SPOP-DAXX interactions, the SPOP assemblies are nearly saturated with DAXX and the collective effect of the DAXX interactions is enough to maintain a condensed, albeit liquid, state. At very high DAXX concentrations there is enough free DAXX to condense into droplets and the SPOP-DAXX assemblies will dissolve in the pure DAXX liquid.

This mechanism shows how the system can switch from a phase driven by the stronger SPOP-DAXX interactions to the weaker DAXX-DAXX interactions by altering the stoichiometry. The small difference in the binding affinities for binding to one vs. two SPOP dimers leads to a narrow range of parameter space where the crosslinked phase is possible (Fig. 2B), which allows for multiple phases to appear over accessible regions of parameter space.

### F. Binding to SPOP increases the DAXX fluid density

Our model can also be used to compute the protein concentrations in the dense phases, which can be compared to the experimental measurements at *c*_*s*_ = 15 *µ*M [23]. At this concentration, the average length of SPOP rods is 50-100 dimers (Fig. 1). This implies a situation where the gel phase has a large number of crosslinks between adjacent rods that will relax very slowly to the equilibrium state of parallel aligned rods. Accordingly, microscope images of the gel phase show a disordered network with locally clustered filaments (Fig. 4 top panel). This is indicative that the SPOP concentration is determined by kinetic arrest. Using the measured SPOP concentration in the gel, *c*_*sg*_ as an input parameter, we can compute the expected concentration of DAXX within the gel. This is given by

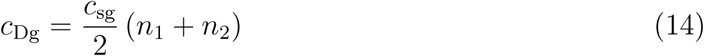

where the factor of two comes from our assumption of cooperative binding to the dimer. Eq. 14 does an excellent job of predicting the DAXX concentration in the gel phase (Fig. 5, left of dashed line), which provides support for our assumption that pairs of SPOP recognition motifs bind cooperatively to SPOP dimers.

**FIG. 5:**
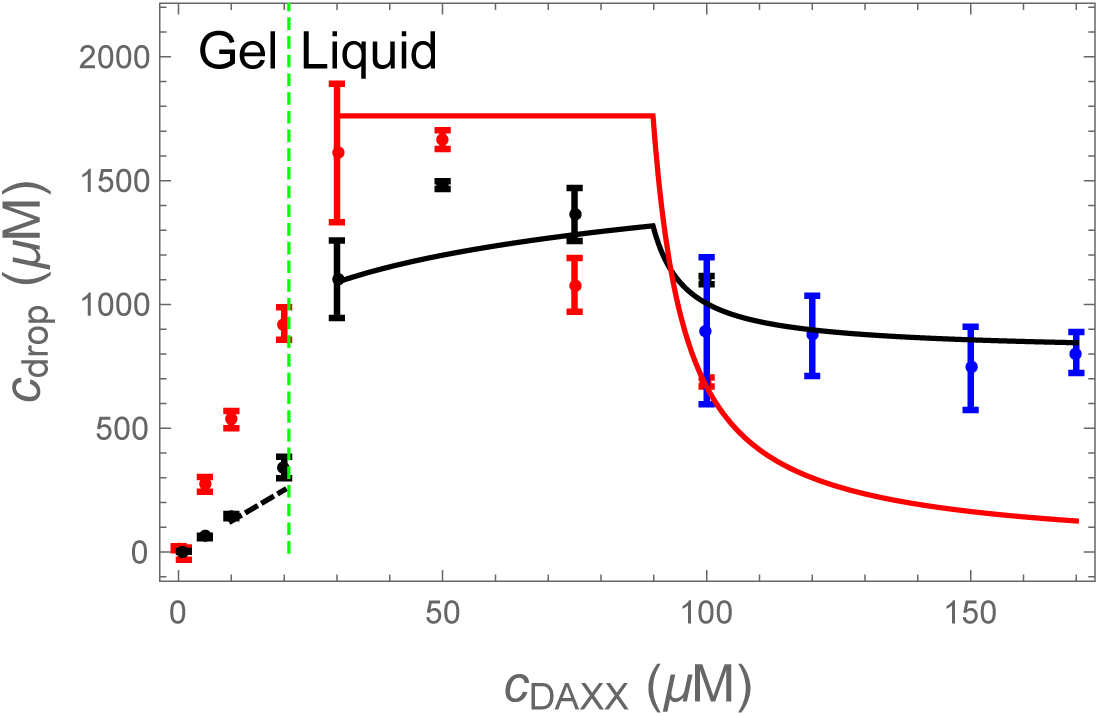
Experimental [23] (points) and computed (lines) protein concentrations in the condensed phases. In the gel phase (left of the dashed line) the SPOP concentration is determined by a kinetic arrest of the network condensation. The cDAXX concentration (black) is given by the binding of cDAXX to this network (Eq. 14). In the liquid phase the SPOP (red) and cDAXX (black) concentrations (Eqs. 32 and 33) are initially given by the excluded volume of the cylindrical SPOP-cDAXX assemblies (plateau region), before relaxing toward the values expected for a pure cDAXX solution (blue points).

As the DAXX concentration increases, the number of crosslinks declines and the system transitions to the liquid state. In the fluid state both SPOP and DAXX concentrations relax to their equilibrium values. These concentrations can be computed from the fraction of SPOP dimers occupied by DAXX molecules and the volume occupied by the brush-like SPOP-DAXX assembly as follows. At the onset of the liquid phase, there is very little free DAXX in the droplets so the SPOP concentration is given by 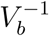, where *V*_*b*_ is the effective volume occupied by a SPOP dimer and the DAXX molecule bound to it. For a long SPOP rod, this volume is a disk with a thickness equal to the dimer diameter and a radius equal to the radius of gyration of DAXX. For the radius of gyration we use ∼ 5 nm based on SAXS measurements (Fig. 6) and random coil estimates for the cDAXX length of 245 amino acids. If a SPOP monomer has a radius ∼ 3 nm, this gives 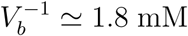, in good agreement with the maximum SPOP concentration in the droplet (Fig. 5). The cDAXX concentration at the fluid onset is 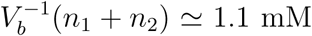, also in good agreement with experiments. This is greater than the 800 *µ*M concentration of pure cDAXX droplets [23]. At greater DAXX concentrations, free DAXX will mix with the droplets. This will dilute the SPOP-DAXX assemblies, causing the SPOP concentration to drop steeply, while the DAXX concentration relaxes toward that of a pure DAXX fluid.

**FIG. 6:**
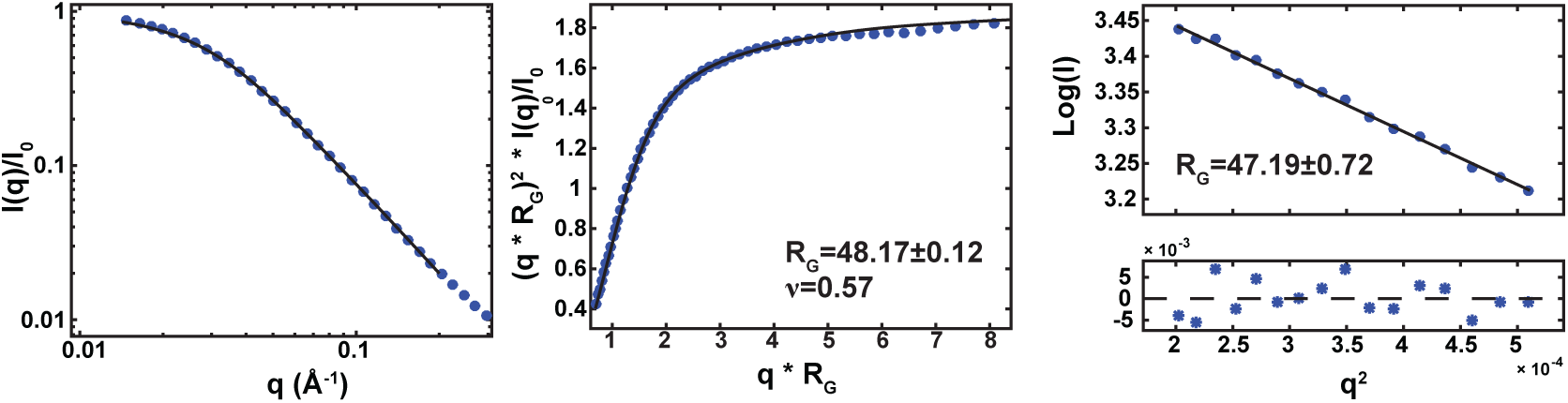
SAXS characterization of cDAXX. (A) Raw SAXS data of cDAXX, where *I*(*q*) normalized by the forward scattering, is plotted versus *q*, defined as *q* = 4*π* sin(2*θ*)*/λ*. Here, *θ* is the scattering angle and *λ* is the X-ray wavelength (∼ 0.1 nm). Experimental data were logarithmically smoothed. Calculated scattering profiles from the empirically derived molecular form factor (MFF) [31] (bottom) are overlaid as a solid line. (B) Raw SAXS data in normalized Kratky representation, logarithmically smoothed. *R*_*g*_ and *n* are a result from the fit to the empirical MFF (solid line). (C) Guinier transformation of the SAXS data. The black line is a linear fit to the Guinier equation with resulting *R*_*g*_ value; the residuals are shown below.

In the Methods section we derive formulas for the relative contributions of the pure DAXX fluid and the SPOP-DAXX brushes. These are compared to the experimental concentrations in Fig. 5. In the liquid phase (right of the dashed line) Eqs. 32 and 33 describe a brush-dominated regime followed by a relaxation toward a pure DAXX fluid as more DAXX is added. This transition is somewhat more gradual than described by the theory indicating the recruitment of additional DAXX molecules to the droplet. This discrepancy is expected due to our neglect of DAXX-DAXX interactions in the assembly free energy (Eq. 4).

## II. DISCUSSION

The SPOP-DAXX system has qualitative similarities to FUS condensates [10] in that both systems form liquid droplets driven by interactions between intrinsically disordered proteins. However, the addition of the second component (SPOP) creates important differences in the phase behavior. First, SPOP provides a backbone that allows DAXX to effectively polymerize into a higher molecular weight entity. This increases the number of favorable contacts that can be formed and shifts the phase boundaries to lower DAXX concentrations. Secondly, the lack of a flexible linker between SPOP dimerization domains permits the formation of solid-like structures in which the movement of the rigid SPOP rods is very slow due to the large number of crosslinkers connecting them. The competing liquid and gel states serve as a kinetic switch, in which the system can toggle between slow and fast dynamics as a function of the molecule concentrations. We speculate that these two states may have implications for decisions to either sequester substrates in gel-like assemblies or turning them over in dense liquids.

Homeostasis requirements place severe restrictions on how living cells can control phase behavior. The system studied here shows several mechanisms that can be used instead of adjusting conditions like temperature, salt concentration, or the protein sequence. First, the effective valency can be controlled by the concentration of the molecules. Increasing the SPOP concentration leads to the assembly of larger scaffolds, while changing the DAXX concentration adjusts the number of crosslinking interactions. Secondly, the gel phase is only possible at intermediate DAXX concentrations, which makes it possible to select the phase assembled by the weaker DAXX-DAXX interactions by saturating the stronger SPOP-DAXX binding. Both phases leverage the effects of multivalency so that relatively weak interactions are sufficient to drive phase separation. However, this can leave the system prone to mutational effects because small perturbations are similarly amplified.

In conclusion, our results highlight the role of the protein architecture in determining the structure of protein assemblies. Furthermore, switches between different types of assemblies are possible from changes in expression levels. We anticipate that network structures contribute to the specificity of condensate assembly and partitioning in cells.

## III. ONLINE METHODS

### A. Calculation of DAXX entropy

To calculate the number of ways to arrange SPOP-DAXX bonds, we start by placing the first site for each of the double-bound DAXX molecules. The number of ways to do this is *M* !*/*(*N*_2_!(*M - N*_2_)!). Next, we place the second binding site for each of these molecules. These sites are more restricted because they are constrained by the first bond. There are *z* possible sites within reach of each DAXX, however, we need to account for the probability that these sites are already occupied. For the placement of the first double attachment, *N*_2_ out of *M* sites are already occupied, so the probability that each of the *z* sites is available is (*M - N*_2_)*/M*. For the placement of the second free tail the probability the neighboring sites are free is (*M -* (*N*_2_ + 1))*/M*. Therefore, the number of ways of placing the second binding sites is

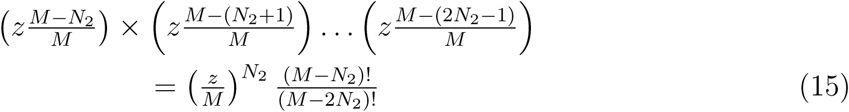

Finally, we need to place the *N*_1_ singly bound DAXX molecules among the remaining *M -* 2*N*_2_ sites leaving *N*_0_ sites unbound. The number of ways to do this is (*M -* 2*N*_2_)!*/*(*N*_0_!*N*_1_!). So the total number of ways to arrange the DAXX molecules is

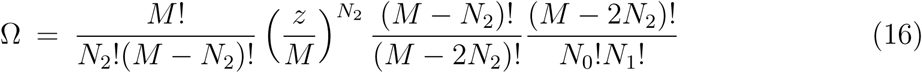

which simplifies to Eq. 5.

### B. Minimization of the gel free energy

With the above expression for the DAXX placement combinatorics and application of Stirling’s approximation, the free energy (Eq. 4) can be written

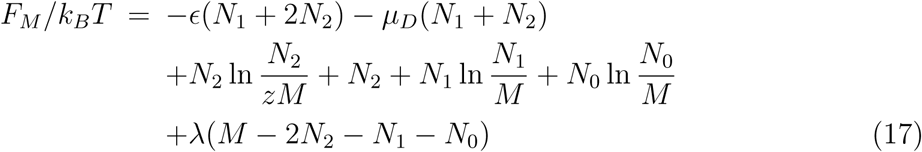

where *λ* is a Lagrange multiplier that will be used to constrain the total number of sites. This can be re-written as

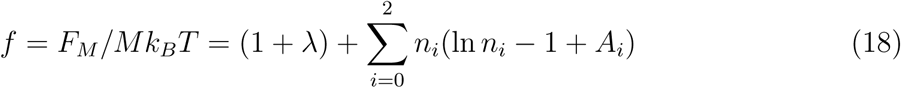

Where *n*_*i*_ = *N*_*i*_*/M* and

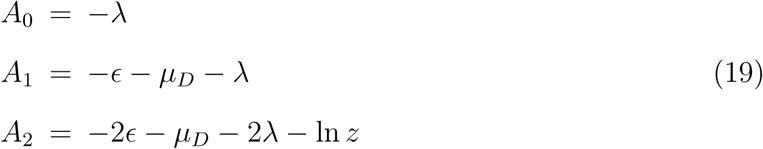

Minimizing the free energy with respect to the site occupancies yields 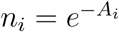, or

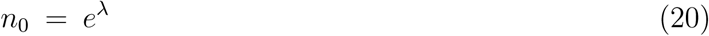

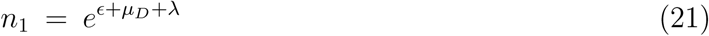

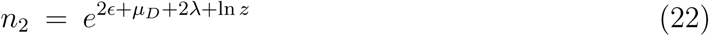

which can be expressed in terms of the concentration of unbound DAXX using the relation *µ*_*D*_ = ln *c*_1*D*_

Inserting Eqs. 20-22 back into Eq. 18 yields Eq. 6. Using Eqs. 8 and 9 with the condition for the total number of sites, 1 = *n*_0_ + *n*_1_ + 2*n*_2_, gives a quadratic equation for *n*_0_ which yields Eq. 7 after selecting the root that gives *n*_0_ → 1 as *c*_1*D*_ → 0.

### C. Calculation of phase boundaries

The vapor-liquid phase boundary is determined from *µ*_*ℓ*_ = *µ*_*v*_. The quantities *c*_*V*_ and *c*_*L*_ appearing in Eqs. 11 and 12 refer to the concentration of SPOP filaments in the vapor and liquid states. We would like to relate these quantities to experiments reporting the concentration of DAXX in the liquid phase. To do this we introduce the quantities *c*_*DV*_ and *c*_*Dℓ*_ representing the concentration of bound DAXX in the vapor and liquid phases, respectively. These are related to the assembly concentrations by *c*_*DV*_ = *N*_*D*_*c*_*V*_ and *c*_*DL*_ = *N*_*D*_*c*_*L*_, where *N*_*D*_ = *n*_1_*L* is the number of single-bound DAXX attached to a SPOP rod. Equating the chemical potentials we have ln(*c*_*DV*_ */c*_0_*N*_*D*_) = ln(*c*_*Dℓ*_*/c*_0_*N*_*D*_) + *ϵ*_*D*_*N*_*D*_ or

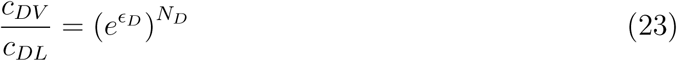

The quantity in parentheses can be estimated from the phase separation of pure DAXX, in which case *N*_*D*_ = 1. In the presence of ficoll the saturation concentration, 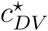, can be constrained by the absence of droplets at 75 *µ*M and their appearance at 100 *µ*M. The resulting droplets have a concentration *c*_*DL*_ = 800 *µ*M [23], therefore, 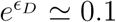. Using Eq. 1 for *c*_*V*_, the phase boundary is given by

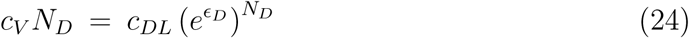

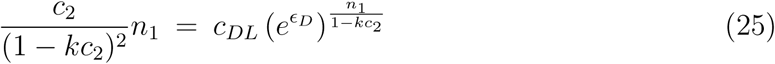

The vapor-gel boundary is given by *µ*_*v*_ = *µ*_*g*_ which yields

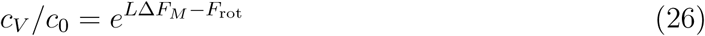

The reference concentration and rotational entropy can be combined into a single parameter, 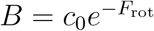, which gives

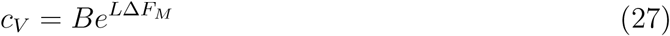

The final phase boundary is obtained from *µ*_*ℓ*_ = *µ*_*g*_. Equating these chemical potentials we have

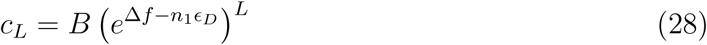

Equations 25, 27, and 28 are solved numerically to determine the phase boundaries. To do this, we require the concentrations of unbound SPOP and DAXX, which are obtained from Eq. 2 and *c*_1*D*_ = *c*_*D*_ *− c*_*s*_(*n*_1_ + *n*_2_), respectively.

### D. Experimental Methods

#### Protein Purification

His-SUMO-SPOP^28−359^ (WT) and His-MBP-SPOP^28−337^ (ΔBACK) were expressed in auto-induction media [32] and purified by Ni NTA affinity chromatography, TEV cleavage, another Ni NTA affinity chromatography (ion exchange for ΔBACK), and SEC chromatography as described before [24, 25]. His-cDAXX^495−740^ was expressed in LB medium and purified by Ni, ion exchange, and SEC chromatography as previously described [23]. For SAXS measurements, the His-tag was cleaved with TEV protease at 4^°C^ overnight before SEC. WT SPOP^28−359^ was labeled with Oregon Green 488 Carboxylic Acid, Succinimidyl Ester, and His-cDAXX^495−740^ was labeled with Rhodamine Red C2 maleimide, as previously described [23].

#### Micrograph images

Samples were prepared with the designated concentrations of proteins, spiked with 0.5 *µ*M fluorescently labeled protein as indicated, and 4% Ficoll 70. Coverslips were prepared and imaged as described before [23] with a Nikon C2 laser scanning confocal microscope at 20x magnification on a (0.8NA) Plan Apo objective, using the same camera settings. Images were processed using Nikon NIS Elements software.

#### Measurement of binding affinities

All direct binding fluorescence anisotropy experiments were performed in 20 mM Tris pH = 7.6, 150 mM NaCl, 5 mM DTT, and 0.01% Triton X-100. Each SPOP construct was serial diluted 10-15 times (from 100 *µ*M to 0.05 *µ*M) into a 384-well plate containing 40 nM Rhodamine labeled His-cDAXX, and 4% Ficoll 70 when indicated. Anisotropy was measured using a CLARIOstar plate reader (BMG LABTECH). *K*_*D*_ values were determined by fitting data points to the fraction bound equation for direct binding given by Roehrl et al. 2004 [33], as described previously [23].

#### SAXS

DAXX samples were prepared for SAXS measurements in a buffer containing 50mM Tris pH 8, 100 mM NaCl, 10 mM DTT and 2 mM TCEP. The buffer was exchanged over a Superdex 75 16/600 column (GE Life Sciences). Buffer was collected 1 column volume after elution for use as a buffer blank. Data was recorded in high-throughput via the mail-in program at the SIBYLS beamline 12.3.1 at the Advanced Light Source at Lawrence Berkeley National Laboratory [34]. Data was recorded on a series of three dilutions (450, 225, 112.5 M) with matched buffer recorded before and after the dilution series. Data included 33 1 second exposures for each sample. All data that was determined to be statistically similar to the first exposure was averaged. For buffer, all 33 exposures were used. For DAXX samples, 12 exposures were used at the highest concentration and 20 exposures were used for the two lower concentrations. Buffer samples before and after the dilution series were identical and were averaged before subtraction. The peak at ∼ 0.39 Å^−1^ associated with the Kapton window was used as a subtraction control. The two lower concentration samples perfectly overlaid and the data was merged to create the final scattering profile. Buffer subtraction, Guinier analysis and Kratky transformation was performed in Matlab (Mathworks). Final data was also fit to an empirically derived molecular form factor for unfolded proteins [31].

### E. Calculation of fluid density

We model the system as having three components, i) a fluid of brush-like SPOP-DAXX assemblies occupying a volume fraction *ϕ*_brush_, ii) a pure DAXX fluid occupying *ϕ*_DAXX_, and a vapor phase occupying *ϕ*_vapor_ = 1 *− ϕ*_brush_ *− ϕ*_DAXX_. The DAXX and brush phases are miscible but we treat them as separate for the moment.

Since the liquid phase is well removed from the vapor phase by the intervening gel phase, it is a good approximation to neglect the SPOP remaining in the vapor phase. Therefore the volume fraction occupied by the SPOP brushes is

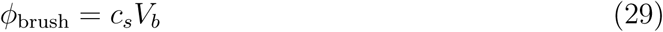

The DAXX liquid volume fraction will be zero as long as the unbound DAXX concentration *c*_1D_ is less than the saturation concentration for droplet formation 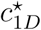. Once the unbound DAXX concentration exceeds this value, it will begin to accumulate in droplets with concentration *c*_*DL*_ such that the remaining solution remains at a fixed concentration 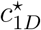. This means that the unbound DAXX must satisfy 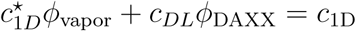 which gives

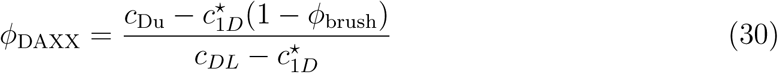

The amount of unbound DAXX is given by

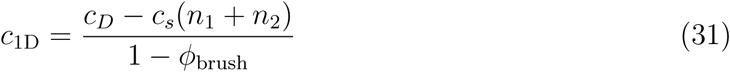

Since the brush liquid and the DAXX liquid are miscible, the protein concentration in the droplets is the weighted average of the two liquids

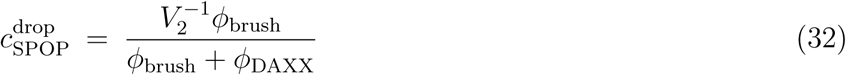

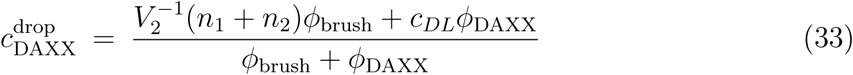

## Acknowledgments

JDS was supported by NIH Grant R01GM107487. T.M. acknowledges funding by NIH grant R01GM112846, St. Jude Childrens Research Hospital, and the American Lebanese Syrian Associated Charities. We thank Melissa Marzahn for purifying SPOP mutants. Microscopy images were acquired at the St. Jude Cell & Tissue Imaging Center, which is supported by SJCRH and NCI (grant P30 CA021765). We also thank Victoria Frohlich and Jennifer Peters for technical help with microscopy. SAXS work was conducted at the Advanced Light Source (ALS), a national user facility operated by Lawrence Berkeley National Laboratory on behalf of the Department of Energy, Office of Basic Energy Sciences, through the Integrated Diffraction Analysis Technologies (IDAT) program, supported by DOE Office of Biological and Environmental Research. Additional support comes from the National Institute of Health project ALS-ENABLE (P30 GM124169) and a High-End Instrumentation Grant S10OD018483.

## Author contributions

JDS developed the theory and performed the calculations. TM conceived of experiments. JDS and TM provided funding. JJB performed purification, binding affinity, and microscopy experiments. EWM purified protein and performed the SAXS measurements. JDS, JJB, and TM wrote the paper.

